# Identification of methylation markers for age and Bovine Respiratory Disease in dairy cattle

**DOI:** 10.1101/2023.12.18.572169

**Authors:** E. Attree, B. Griffiths, K. Panchal, D. Xia, D. Werling, G. Banos, G. Oikonomou, A. Psifidi

## Abstract

Methylation profiles of animals is known to differ by age and disease status. Bovine respiratory disease (BRD), a complex infectious disease, primarily affects calves and has significant impact on animal welfare and the cattle industry, predominantly from production losses. BRD susceptibility is multifactorial, influenced by both environmental and genetic factors. We investigated the epigenetic profile of BRD susceptibility in calves and age-related methylation differences between healthy calves and adult dairy cows using Reduced Representation Bisulfite Sequencing (RRBS).

We identified 3,452 genes within differentially methylated regions between calves and adults. Functional analysis revealed enrichment of developmental pathways including cell fate commitment and tissue morphogenesis. Between healthy and BRD affected calves, 964 genes were identified within differentially methylated regions. Immune and vasculature regulatory pathways were enriched and key candidates in BRD susceptibility involved in complement cascade regulation, vasoconstriction and respiratory cilia structure and function were identified.

## Introduction

Epigenetic changes are genetic modifications that influence gene regulation, without changing the DNA sequence. DNA methylation plays an important role in the development of disease [1, 2] and ageing [3–6]. Although there are a lot of relevant studies in humans and mice, there are limited studies in cattle. Differences in DNA methylation caused by environmental factors alter transcriptional regulation, for example by silencing genes, and thus, differential methylation is an important epigenetic modification that can greatly impact development and disease. DNA methylation changes are associated with age in humans and animals [7, 8], these known age-related changes can be used to construct epigenetic clocks. Epigenetic clocks are tools in the prediction of genetic age of tissues and individuals that can be used to predict chronological age or assist in identification of the influence of biological differences such as disease, individual genetics and environmental factors [4]. Epigenetic clocks have been constructed for human and mouse tissues, using methylation data from multiple tissue types [4, 6]. In cattle, only very recently epigenetic clocks have been constructed to predict chronological age [3, 9] and for the prediction of oocyst epigenetic age and therefore reproductive aging, an important consideration in the dairy industry and potentially a useful model for human reproductive aging [10].Moreover, the role of DNA methylation in dairy cattle mastitis caused by *Staphylococcus aureus* [11] and *Escherichia coli* [12] has been investigated.

In addition to mastitis, early life calf losses cause a significant economic drain on the dairy industry as a dairy cow does not generate direct income until after its first calving [13]. The cost of rearing a replacement dairy cow has previously been estimated as 15-20% of the animal’s economic production value [14]. Those heifers that die early or have to be culled before the end of their first lactation do not reimburse the economic cost of their rearing. Early life diseases have a significant impact on animal welfare, and thus on the cattle industry as a whole. Losses are typically attributed to impaired growth and consequential reduction in carcass value, the cost of disease management and treatment and, in severe cases, mortality [15–18].

Excluding abortions, still-births and calves that die within the first 24 hours after delivery, the incidence of neonatal mortality (calves aged from 1-28 days) has been reported to account for up to 12% [19, 20]. The two most prevalent diseases contributing to this include diarrhoea and bovine respiratory disease complex (BRDC) [21] with clinical signs observed in pre weaned calves at a prevalence of 13.8-21.6% and 8.1-22.8% respectively [22–24].

Multiple pathogens have been reported to contribute to the development of BRDC, including bacteria such as *Histophilus somni*, *Mannheimia hemolytica, Mycoplasma bovis* and *Pasteurella multocida* [25–30] and viruses such as bovine coronavirus (BoCoV, bovine viral diarrhoea virus (BVDV), bovine herpesvirus-1 (BHV-1), bovine reovirus, bovine respiratory syncytial virus (BRSV) and parainfluenxavirus-3 (BoPi-3) [18, 31–34]. Newborn animals are at increased risk of infection with these pathogens due to their not yet fully developed adaptive immune system, environmental stressors (mainly weaning, transportation and contact with animals from other source) that increase potentially immunosuppression [35]. Management of BRDC is typically by either curative of prophylactic administration of antibiotics [18, 36]. This aids potentially the development of antimicrobial resistance [18] and therefore poses a risk to both, animal and human health.

An animal’s response to complex diseases such as BRD is multifactorial, influenced by both the environment, the pathogen and individual genetic profiles. Previous studies have investigated the heritability of resistance to BRDC with estimates ranging from 0.07 to 0.29 [37–39]. Moreover, it has been suggested that an epigenetic component may be involved since disease episodes early in a calf’s life has been reported to negatively impact lifetime performance and age at first calving [40, 41]. Therefore, studying the methylation profile of these calves may provide further data for future breeding programs designed to improve herd resistance [37, 38, 42, 43] as well as increase our understanding of the underlying mechanisms of disease susceptibility. This endeavour would be greatly advantageous in improving genetic resistance, welfare of animals and subsequently reducing resultant economic losses. Further, reducing the instance of antibiotic treatment of non-affected animals would consequently reduce the risk of the development of antimicrobial drug resistance. In the present study we examined differential methylation between healthy calves and cows to establish baseline age related differential methylation for the purpose of identification of potential markers of genetic age. We also examined differences in methylation with respect to health status of calves to identify differential methylation between healthy and BRD affected that may convey resistance to disease for the purpose of identifying potential targets for genetic improvement of dairy cattle.

## Results

The health of 14 calves was assessed over an eight-week period with Wisconsin scores [44] recorded at weeks one, five and eight, table 1. Calves were ranked best to worst based on total score. Six of these calves were selected to compare their blood methylation profiles, three calves with the lowest total score were taken forward as the ‘healthy’ samples and three calves with some of the worst scores (scores 10-12 out of 14) were taken forward as the ‘diseased’ calf samples. Genetic relatedness of the calves was considered during selection: selected cases and controls were half siblings with calves 3 and 12 being twins (full siblings) with different BRD diagnoses (Supplementary table S1).

**Table 1:** Wisconsin scores for weeks 1, 5 and 8 of the study and the total score over the eight weeks, the number of times respiratory disease was observed over the eight-week period and for each calf the lung scanning score as measured at week 8.

| **Calf** | **Week 1**<br>**Wisconsin**<br>**score total** | **Week 5**<br>**Wisconsin**<br>**score total** | **Week 8**<br>**Wisconsin**<br>**score total** | **Respiratory**<br>**Disease**<br>**Frequency\*** | **Lung**<br>**Scanning**<br>**Score** |
| --- | --- | --- | --- | --- | --- |
| 1 | 0 | 0 | 0 | 0 | 0 |
| 2 | 0 | 0 | 0 | 0 | 0 |
| 3 | 1 | 0 | 1 | 0 | 1 |
| 10 | 0 | 1 | 1 | 1 | 2 |
| 11 | 0 | 3 | 3 | 1 | 0 |
| 12 | 4 | 0 | 3 | 1 | 2 |

\*The number of times respiratory disease was detected from the week 1, 5 and 8 checks.

Two adult cows included in the analysis were also from the same farm as the calves, ensuring that all animals were raised under the same management,

### Comparison of methylation profiles between healthy calves and cows

Firstly, the methylation profiles of healthy calves and cows were compared to form a baseline understanding of methylation differences based on animal age. Clear clustering and separation between the methylation profiles of cows and calves was observed (Figure 1A and B). Between adults and calves 2177 (0.89%) bases were hypermethylated and 2801 (1.16%) bases were hypomethylated.

**Figure 1:**
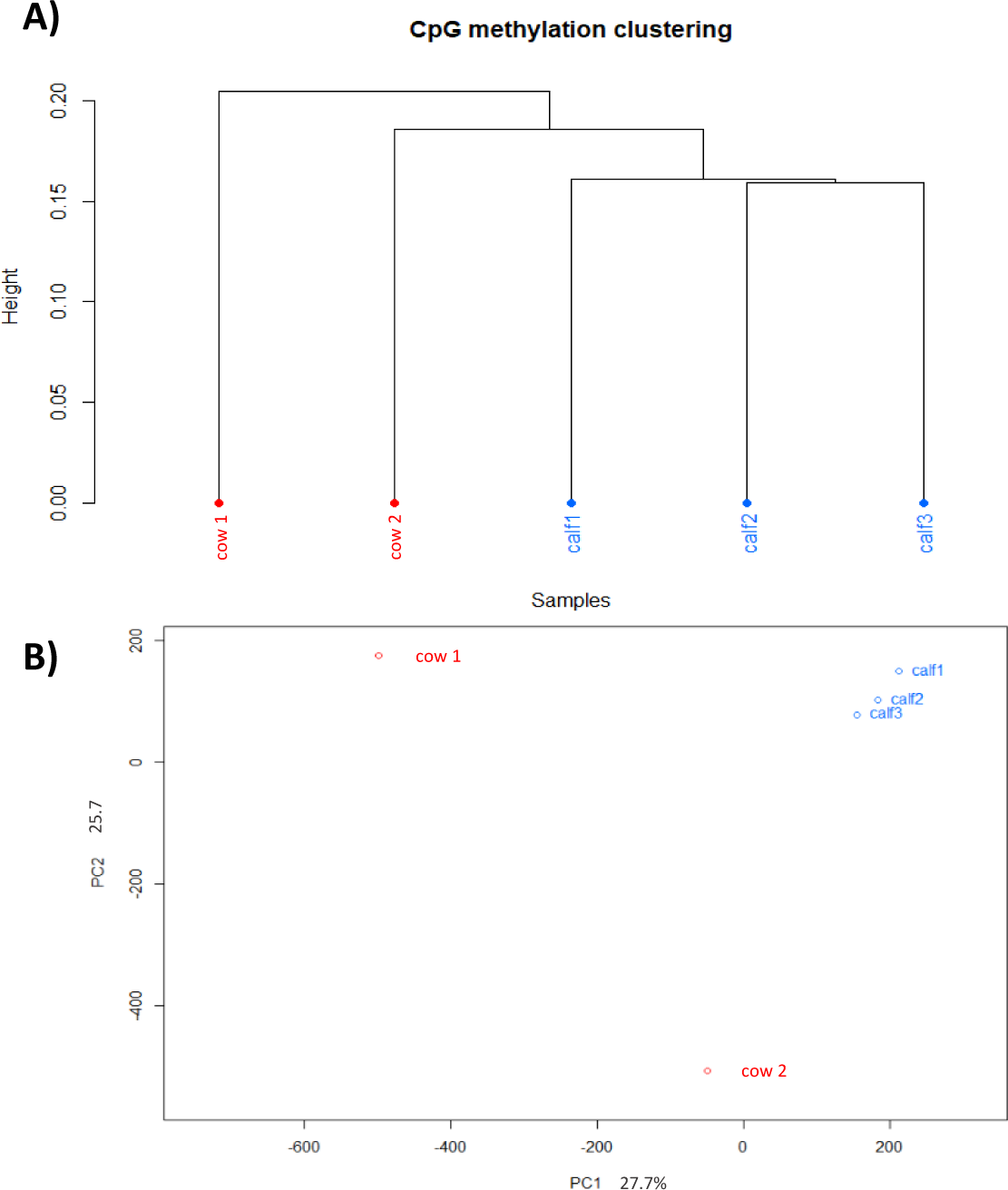
A) CpG methylation clustering of healthy calves and cows displayed by dendrogram and B) PCA analysis indicating the clustering and separation of cows and calves.

A total of 4713 regions were differentially methylated (≥25% difference of methylation between groups, p ≤0.05 and q ≤0.01) at the time of sampling between cow and calf samples, corresponding to 3452 genes. The distribution of differential methylation after annotation of gene features indicated that 6% of differential methylation was identified in promoter regions, 10% in exon, 30% in intronic and 54% in intergenic regions.

Functional enrichment analysis by Database for Annotation, Visualization and Integrated Discovery (DAVID) [45, 46] was conducted for all genes identified with statistically differentially methylated regions. The biological process terms with the greatest fold enrichment (>6) were: midbrain-hindbrain boundary morphogenesis (GO:0021555), protein farnesylation (GO:0018343), fibroblast apoptotic process (GO:0044346), atrioventricular bundle cell differentiation (GO:0003167), negative regulation of dendritic spine development (GO:0061000), actin-mediated cell contraction (GO:0070252), establishment of planar polarity of embryonic epithelium (GO:0042249), Wnt signaling pathway involved in somitogenesis (GO:0090244), regulation of glomerular filtration (GO:0003093), tube formation (GO:0035148) and negative regulation of vascular smooth muscle cell differentiation (GO:1905064). No terms were, however, found to be statistically significant (>0.10) after Bonferroni correction. In functional annotation clustering analysis, 37 clusters with enrichment scores >1, four with enrichment scores >3 were identified. The clusters with the greatest enrichment scores included: DNA-templated transcription, initiation (enrichment score 5.12), Protein kinase, catalytic domain (enrichment score 5.09), Src homology-3 domain (enrichment score 3.99) and Pleckstrin homology domain (enrichment score 3.93). Panther overrepresentation test found 15 statistically significant (p≤0.05 and FDR≤0.05) under-represented GO biological process’ primarily involved in cellular and metabolic process’ and Reactome pathway analysis identified eight significantly enriched pathways (p≤0.05 and FDR≤0.05) involved in signal transduction, interactions at synapses and the neuronal system.

Age related methylation differences have previously been reported positionally within 50kb of the TSS [47], this threshold has also been previously applied to define *cis* regions of differential methylation [48]. Functional analysis of identified differential methylation was, therefore, restricted to 50kb either side of the TSS, resulting in 2553 genes. Functional enrichment analysis by DAVID found the terms with the greatest fold enrichment (>10) included: atrioventricular bundle cell differentiation (GO:0003167), regulation of glomerular filtration (GO:0003093) and establishment of planar polarity of embryonic epithelium (GO:0042249). These terms were not however found to be statistically significant after Bonferroni correction. In functional annotation clustering analysis 34 clusters were identified with enrichment scores >1 with three of those >3: protein kinase domain (enrichment score 3.97), DNA-templated transcription, initiation (enrichment score 3.95) and SH3 domain (enrichment score 3.56). Two Reactome pathways were identified as significantly enriched (p ≤0.01, FDR ≤0.05): O-linked glycosylation (fold enrichment 2.85) and Signalling by Receptor Tyrosine Kinases (fold enrichment 1.78).

GO enrichment analysis of these genes by PANTHER Overrepresentation Test identified 25 over and under enriched terms (p ≤0.0001, FDR ≤0.05) with the greatest enrichment observed in the regulation of growth, for example: regulation of growth, cell migration, cell motility, regulation of cell communication and positive regulation of signalling, presented in Table 2 and Figure 2. GO terms from the Panther overrepresentation test were further simplified and visualised using REVIGO to identify the key enriched pathways, presented in Figure 3. We found enrichment for organ morphogenesis, cell adhesion, adult behaviour, osteoblast differentiation growth hormone receptor signalling pathway and developmental pathways including Wnt signalling (an essential evolutionary conserved pathway involved in embryonic organogenesis, cell fate, motility and neural organisation [49]) and the ERK1/2 cascade (involved in cell proliferation, differentiation, apoptosis and stress responses [50, 51]). Immune related pathways were also found to be enriched, namely B1-B cell homeostasis, the cells initially generated early in an animals life (foetal and early neonatal) [52, 53] and the B cell subgroup reportedly associated with the generation of an antibody response during infection and vaccination [54].

**Table 2:** The 25 over and under enriched GO biological process terms as identified by PANTHER Overrepresentation Test of all genes identified with differentially methylated regions within 50kb of the TSS.

| GO biological process | fold Enrichment | P value | FDR |
| --- | --- | --- | --- |
| regulation of growth (GO:0040008) | 1.76 | 5.5E-05 | 3.6E-02 |
| cell migration (GO:0016477) | 1.55 | 3.0E-05 | 2.5E-02 |
| cell motility (GO:0048870) | 1.5 | 2.4E-05 | 2.3E-02 |
| negative regulation of signal transduction (GO:0009968) | 1.46 | 5.2E-05 | 3.5E-02 |
| negative regulation of RNA biosynthetic process (GO:1902679) | 1.43 | 7.3E-05 | 4.4E-02 |
| negative regulation of DNA-templated transcription (GO:0045892) | 1.43 | 8.7E-05 | 5.0E-02 |
| negative regulation of response to stimulus (GO:0048585) | 1.41 | 3.5E-05 | 2.7E-02 |
| positive regulation of signaling (GO:0023056) | 1.4 | 3.4E-05 | 2.7E-02 |
| positive regulation of cell communication (GO:0010647) | 1.39 | 4.2E-05 | 3.0E-02 |
| regulation of signal transduction (GO:0009966) | 1.36 | 2.4E-07 | 4.2E-04 |
| negative regulation of cellular metabolic process (GO:0031324) | 1.36 | 1.6E-05 | 1.6E-02 |
| regulation of cell communication (GO:0010646) | 1.34 | 1.4E-07 | 3.9E-04 |
| regulation of signaling (GO:0023051) | 1.34 | 2.5E-07 | 3.9E-04 |
| negative regulation of nitrogen compound metabolic process (GO:0051172) | 1.31 | 6.6E-05 | 4.1E-02 |
| regulation of response to stimulus (GO:0048583) | 1.27 | 5.3E-06 | 6.3E-03 |
| negative regulation of cellular process (GO:0048523) | 1.24 | 4.1E-06 | 5.4E-03 |
| negative regulation of biological process (GO:0048519) | 1.22 | 5.8E-06 | 6.4E-03 |
| sensory perception (GO:0007600) | 0.65 | 2.4E-05 | 2.2E-02 |
| sensory perception of chemical stimulus (GO:0007606) | 0.54 | 6.6E-07 | 9.4E-04 |
| detection of stimulus (GO:0051606) | 0.53 | 2.2E-07 | 4.4E-04 |
| detection of stimulus involved in sensory perception (GO:0050906) | 0.52 | 2.1E-07 | 5.1E-04 |
| detection of chemical stimulus involved in sensory perception (GO:0050907) | 0.49 | 7.8E-08 | 3.7E-04 |
| sensory perception of smell (GO:0007608) | 0.49 | 8.3E-08 | 3.0E-04 |
| detection of chemical stimulus (GO:0009593) | 0.48 | 2.9E-08 | 4.1E-04 |
| detection of chemical stimulus involved in sensory perception of smell (GO:0050911) | 0.47 | 3.1E-08 | 2.2E-04 |

**Figure 2:**
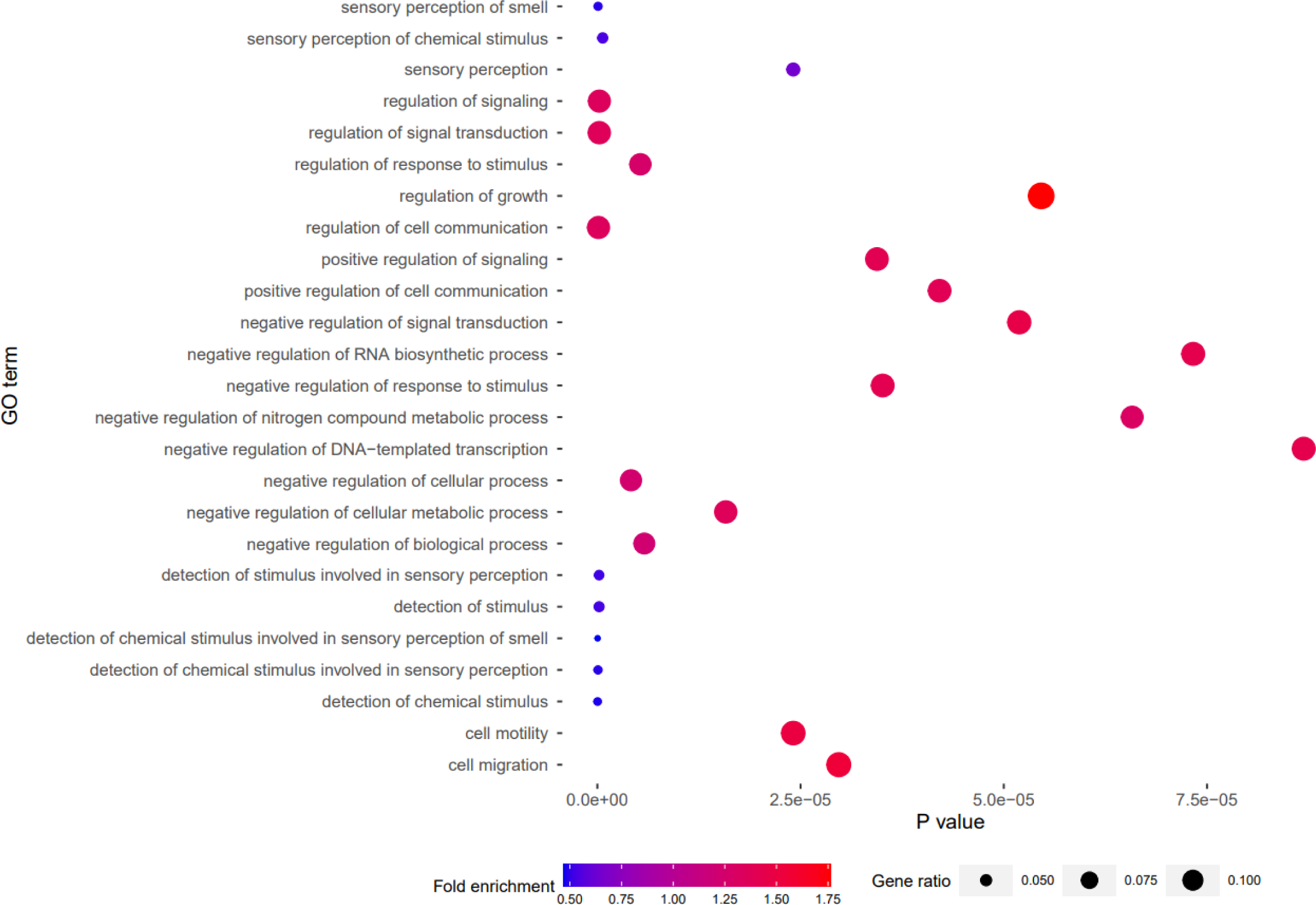
Graphical representation of over and under-represented GO biological process’ identified by PANTHER Overrepresentation Test of the genes identified within 50kb of differentially methylated regions identified between healthy cows (as control) and calves. Gene ratio was calculated by gene count per term / number of genes assigned the term in the genome.

**Figure 3:**
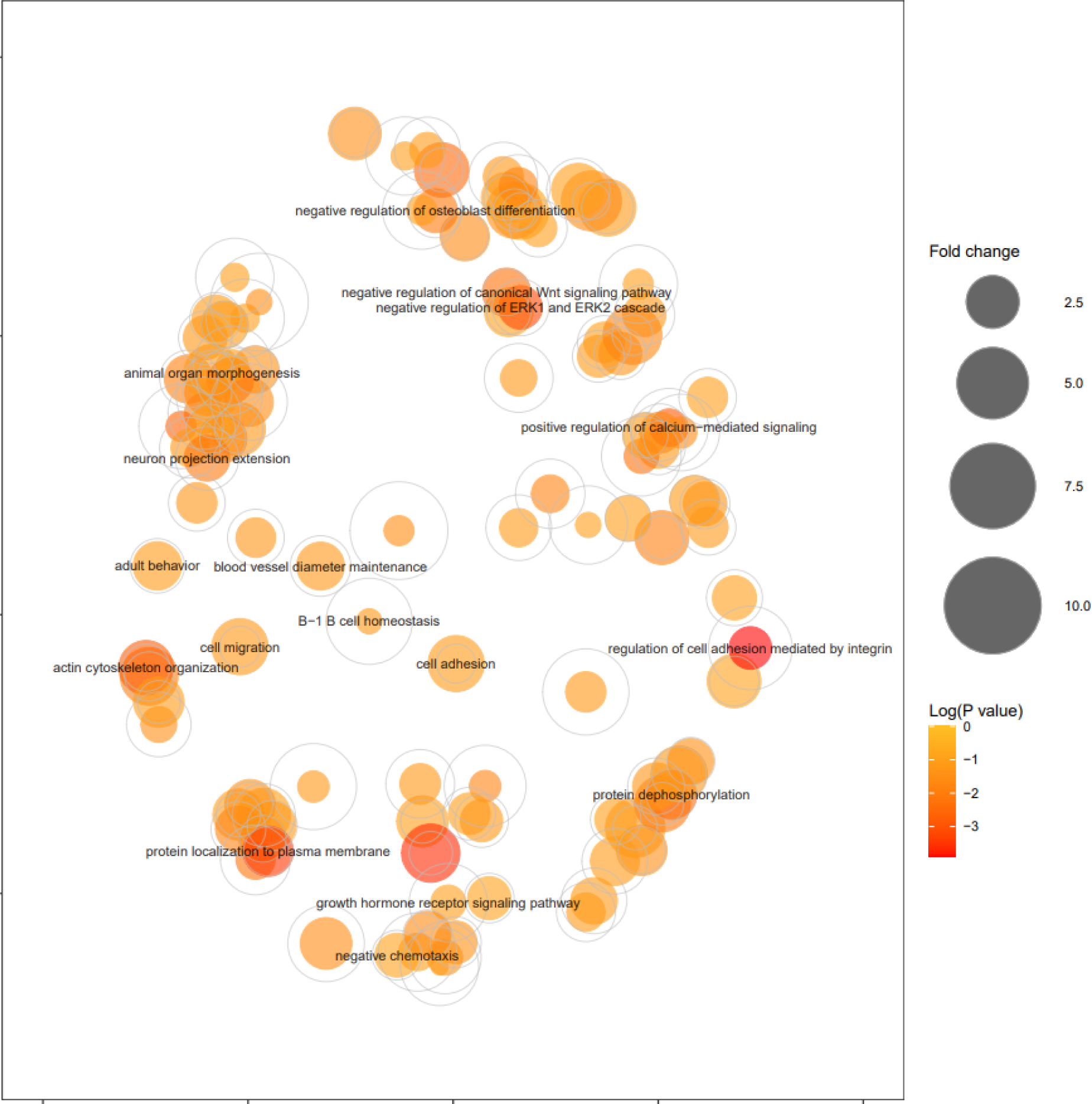
Graphical visualisation by REVIGO of over and under-represented GO biological process’ identified by Panther overrepresentation test of the genes identified within 50kb of differentially methylated regions identified between healthy cows and calves.

Further stringent restriction of region of methylation with respect to the TSS to 10kb either side was applied for further functional enrichment. A single significantly enriched GO biological process term after Bonferroni correction was identified: GO:0006352∼DNA-templated transcription, initiation (7.36 fold enrichment, 4.94E-05 Bonferroni correction). Nine functional annotation clusters were identified with enrichment score >1, one with a score >3: DNA-templated transcription, initiation (enrichment score 5.64), Pleckstrin homology-like domain (enrichment score 2.12), Serine/threonine-protein kinase (enrichment score 1.31), Netrin domain (enrichment score 1.25), negative regulation of endopeptidase activity (enrichment score 1.22), AIG1 (enrichment score 1.19), Longevity regulating pathway (enrichment score 1.17), Ras-association (enrichment score 1.1) and gene silencing by miRNA (enrichment score 1.06). Panther overrepresentation test found six statistically significant under-represented GO biological process terms: cellular process (GO:0009987, fold change 0.72), biological_process (GO:0008150, fold change 0.70), metabolic process (GO:0008152, fold change 0.63), organic substance metabolic process (GO:0071704, fold change 0.59), nitrogen compound metabolic process (GO:0006807, fold change 0.58) and primary metabolic process (GO:0044238, fold change 0.58). A protein interaction network analysis of the genes with differential methylation identified within 10kb either side of the TSS was generated using STRING (version 11.5) [55], presented in Figure 4. Clustering of interacting proteins into three clusters revealed pathways involved in AMPK signalling pathway, intracellular signal transduction, protein kinase, ATP binding transcriptional regulation and MAPK pathway.

**Figure 4:**
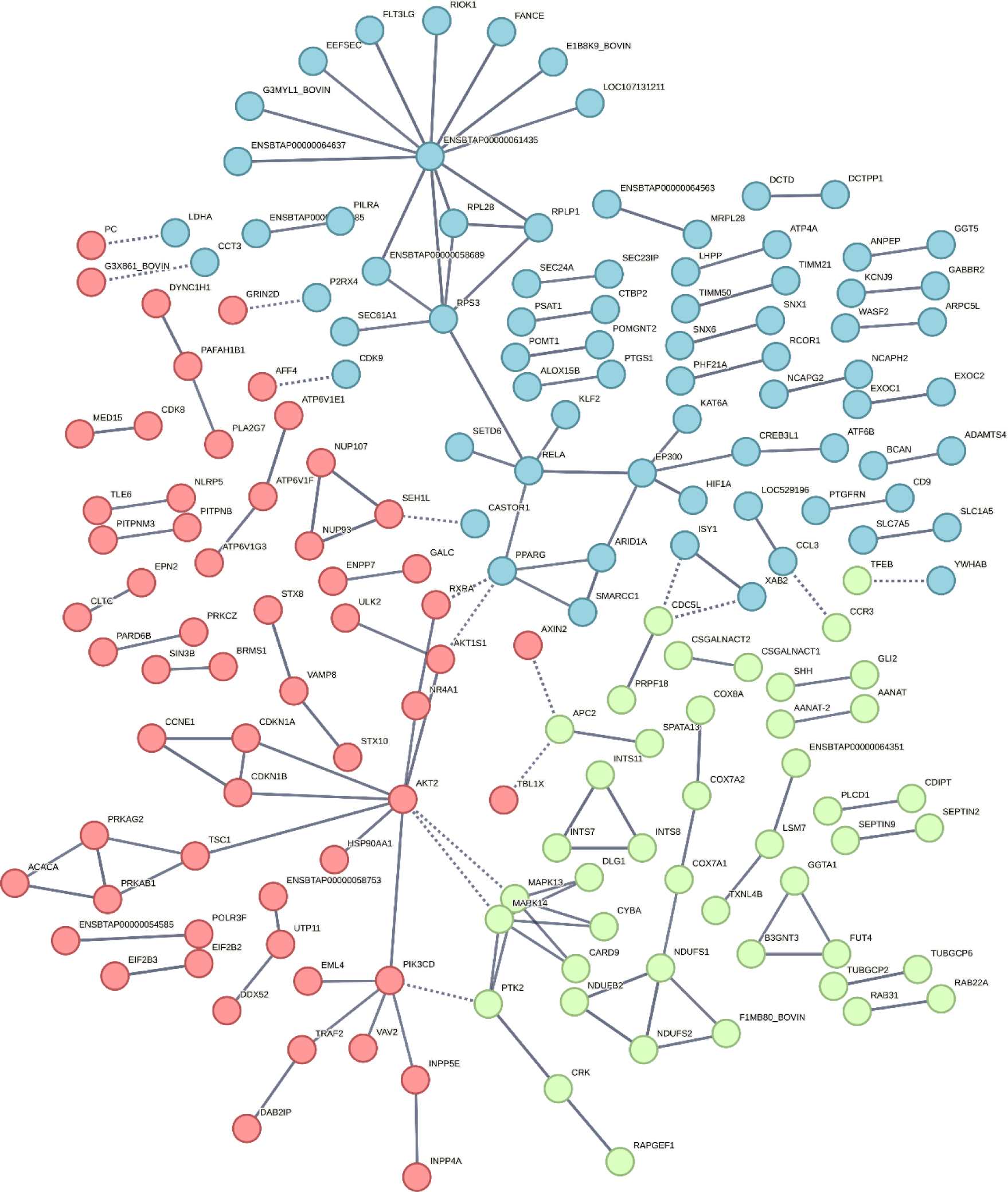
String protein interaction network of identified genes with differential methylation between healthy cows and calves within 10kb of a TSS.

Extension of differential methylation analysis to base level identification of differential methylation in promotor and exon regions identified 1079 genes with differential methylation in their promotor regions and 954 in their exon regions. No functional enrichment was however observed by Panther overrepresentation test for GO biological process’ or Reactome pathways.

### Comparison of methylation profiles between healthy calves and calves diagnosed with BRD

The methylation profiles of healthy and diseased calves, with respect to BRD, were compared in order to identify methylation differences that may be related to resistance to BRD. The analysis of differential methylation between healthy and diseased animals, revealed 1541 regions (0.58%) bases were hypermethylated and 1511 (0.57%) regions were hypomethylated. A total of 1029 regions were differentially methylated (≥25% difference of methylation between groups q ≤0.01) between healthy and diseased samples corresponding to 964 genes. The distribution of differential methylation after annotation of gene features indicated 5% of differential methylation was identified in promoter regions, 9% in exonic regions, 30% in intronic regions and 56% in intergenic regions.

Functional enrichment analysis by DAVID was conducted for all genes with differentially methylated regions. The greatest fold enrichment of biological process (fold change >10) was observed for: regulation of centrosome cycle (GO:0046605, fold change 19.51), positive regulation of renal sodium excretion (GO:0035815, fold change 19.51), positive regulation of plasma cell differentiation (GO:1900100, fold change 19.51), zygotic specification of dorsal/ventral axis (GO:0007352, fold change 19.51), negative regulation of non-canonical Wnt signalling pathway (GO:2000051, fold change 14.63) and positive regulation of T-helper 2 cell cytokine production (GO:2000553, fold change 11.15). No terms were, however, found to be statistically significant after Bonferroni correction. After functional annotation clustering analysis, 25 clusters were identified with enrichment scores >1, those with the highest scores included: cell-cell adhesion via plasma-membrane adhesion molecules (enrichment score 2.32), Fibronectin, type III (enrichment score 2.32) and Basic-leucine zipper domain (enrichment score 1.9).

Panther Overrepresentation test of these genes found 24 statistically significant over (21) and under (3) enriched GO biological process terms (p≤0.05, FDR≤0.05), detailed in Table 3. Functional pathway analysis by Reactome of all genes associated with differentially methylated regions identified three over-represented pathways, including: neuronal system (fold enrichment 2.29, p-value 8.19E-06, FDR 5.42E-03), adaptive immune system (fold enrichment 1.79, p-value 4.55E-05, FDR 1.51E-02) and signal transduction (fold enrichment 1.4, p-value 3.94E-05, FDR 1.74E-02).

**Table 3:** The 24 over and under enriched GO biological process terms as identified by PANTHER Overrepresentation Test of all genes identified with differentially methylated regions.

| GO biological process complete | fold Enrichment | P value | FDR |
| --- | --- | --- | --- |
| cell junction assembly (GO:0034329) | 2.54 | 7.7E-05 | 4.6E-02 |
| cell junction organization (GO:0034330) | 2.18 | 1.6E-05 | 1.9E-02 |
| neuron development (GO:0048666) | 1.94 | 3.6E-06 | 6.5E-03 |
| cell morphogenesis (GO:0000902) | 1.89 | 7.1E-05 | 4.6E-02 |
| neuron differentiation (GO:0030182) | 1.83 | 4.2E-06 | 6.7E-03 |
| generation of neurons (GO:0048699) | 1.82 | 2.1E-06 | 4.9E-03 |
| neurogenesis (GO:0022008) | 1.75 | 2.8E-06 | 5.7E-03 |
| plasma membrane bounded cell projection organization (GO:0120036) | 1.69 | 2.6E-05 | 2.5E-02 |
| cell projection organization (GO:0030030) | 1.67 | 3.3E-05 | 2.6E-02 |
| nervous system development (GO:0007399) | 1.58 | 4.7E-06 | 6.8E-03 |
| system development (GO:0048731) | 1.51 | 6.4E-08 | 9.1E-04 |
| animal organ development (GO:0048513) | 1.50 | 1.0E-06 | 3.7E-03 |
| cell development (GO:0048468) | 1.48 | 4.4E-05 | 3.0E-02 |
| anatomical structure morphogenesis (GO:0009653) | 1.48 | 3.5E-05 | 2.6E-02 |
| multicellular organism development (GO:0007275) | 1.46 | 9.1E-08 | 6.5E-04 |
| negative regulation of cellular metabolic process (GO:0031324) | 1.45 | 7.6E-05 | 4.7E-02 |
| cellular developmental process (GO:0048869) | 1.38 | 1.9E-05 | 2.1E-02 |
| cell differentiation (GO:0030154) | 1.38 | 2.7E-05 | 2.3E-02 |
| anatomical structure development (GO:0048856) | 1.36 | 6.1E-07 | 2.9E-03 |
| developmental process (GO:0032502) | 1.33 | 1.1E-06 | 3.2E-03 |
| cellular component organization (GO:0016043) | 1.27 | 1.3E-05 | 1.6E-02 |
| detection of stimulus (GO:0051606) | 0.48 | 3.5E-05 | 2.5E-02 |
| detection of stimulus involved in sensory perception (GO:0050906) | 0.46 | 2.6E-05 | 2.4E-02 |
| detection of chemical stimulus involved in sensory perception of smell (GO:0050911) | 0.43 | 2.4E-05 | 2.4E-02 |

A 50kb threshold for methylated regions either side of the TSS was applied as a previously defined *cis* region [48] and previously associated with response to bacterial infection in humans [56]. A total of 2553 genes with differentially methylated regions within 50kb (either side) of the TSS were taken forward for functional analysis. Functional enrichment analysis by DAVID found the greatest fold change (>6) in terms involved in angiogenesis and vasculature, osteogenesis and immune regulation. Specifically, these include: aortic valve morphogenesis (GO:0003180), positive regulation of chondrocyte differentiation (GO:0032332), positive regulation of calcineurin-NFAT signalling cascade (GO:0070886), negative regulation of vascular permeability (GO:0043116), positive regulation of T-helper 2 cell cytokine production (GO:2000553). No terms were, however, found statistically significant after Bonferroni correction. After functional annotation clustering 14 clusters were identified with enrichment scores >1. The greatest enrichment was identified for the clusters calcium-dependent cysteine-type endopeptidase activity (enrichment score 2), Fibronectin, type III (enrichment score 1.78) and dilute domain (enrichment score 1.78).

GO terms from the DAVID functional enrichment analyses were further simplified and visualised using REVIGO to identify the key enriched pathways (Figure 5). Those of particular relevance to BRD that were identified included those involved in epithelial cell regulation and vascular permeability and immune regulation, for example: positive regulation of T-helper 2 cell cytokine production (lymphocytes involved in the adaptive immune response recruiting other activated immune cells and the production of antibodies [57]), negative regulation of vascular permeability, angiogenesis and cell adhesion (Figure 5).

**Figure 5:**
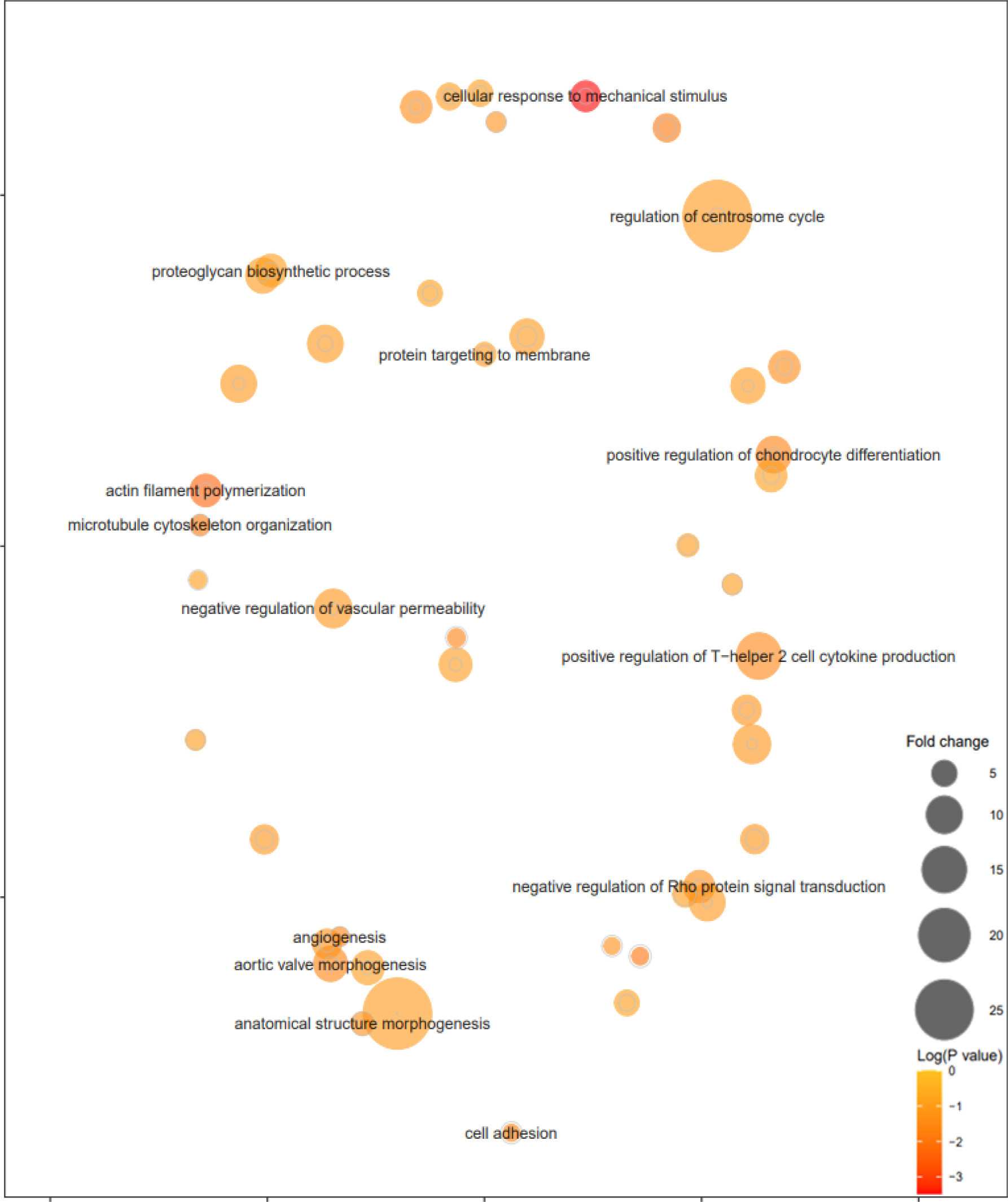
Graphical visualisation by REVIGO of over and under-represented GO biological process’ identified by DAVID functional enrichment analysis of the genes identified within 50kb of differentially methylated regions identified between healthy calves and those diagnosed with BRD.

Panther Overrepresentation test found nine underrepresented GO terms, no overrepresentation was observed, with statistical significance (p≤0.05, FDR≤0.05). These were primarily in metabolic process’: regulation of biological process (GO:0050789), metabolic process (GO:0008152), cellular process (GO:0009987), biological regulation (GO:0065007), biological_process (GO:0008150), organic substance metabolic process (GO:0071704), nitrogen compound metabolic process (GO:0006807), primary metabolic process (GO:0044238) and cellular nitrogen compound metabolic process (GO:0034641).

More stringent restriction of region of methylation with respect to the TSS to 10kb either side was applied for further functional enrichment. The greatest fold enrichment as identified by DAVID functional enrichment analysis was for the GO biological process terms: regulation of centrosome cycle (GO:0046605, fold change 58.34), plasma membrane fusion (GO:0045026, fold change 38.89), DNA catabolic process, exonucleolytic (GO:0000738, fold change 29.17), testosterone biosynthetic process (GO:0061370 fold change 29.17), endosome to lysosome transport via multivesicular body sorting pathway (GO:0032510, fold change 23.34), negative regulation of interleukin-4 production (GO:0032713, fold change 23.34), positive regulation of signal transduction by p53 class mediator (GO:1901798, fold change 19.45), fibroblast migration (GO:0010761, fold change 13.46) and proteoglycan biosynthetic process (GO:0030166, fold change 12.50). Functional annotation clustering revealed six clusters with enrichment scores ≥1: signal transduction (enrichment score 1.43), FAD binding (enrichment score 1.42), stress response (enrichment score 1.09), Pathways of neurodegeneration - multiple diseases (enrichment score 1.05), actin cytoskeleton organization (enrichment score 1.04) and Immunoglobulin domain (enrichment score 1.0). Panther Overrepresentation test identified 15 significantly under-represented GO biological process terms primarily involved in metabolic process’: cellular process (GO:0009987, fold change 0.51), biological_process (GO:0008150, fold change 0.49), regulation of biological process (GO:0050789, fold change 0.46), biological regulation (GO:0065007, fold change 0.46), metabolic process (GO:0008152, fold change 0.42), organic substance metabolic process (GO:0071704, fold change 0.37), macromolecule metabolic process (GO:0043170, fold change 0.33), primary metabolic process (GO:0044238, fold change 0.28), nitrogen compound metabolic process (GO:0006807, fold change 0.25), cellular nitrogen compound metabolic process (GO:0034641, fold change 0.06), cellular aromatic compound metabolic process (GO:0006725, fold change <0.01), nucleic acid metabolic process (GO:0090304, fold change <0.01), nucleobase-containing compound metabolic process (GO:0006139, fold change <0.01), organic cyclic compound metabolic process (GO:1901360, fold change <0.01) and heterocycle metabolic process (GO:0046483, fold change <0.01). No over-represented terms were identified. A protein interaction network analysis of genes with differential methylation identified within 10kb of a TSS was generated using STRING (version 11.5) [55], Figure 6. Protein interaction networks identified revealed enrichment of pathways involved in the mitochondrial membrane respiratory chain, Cytochrome P450 pathway, ribosomal activity, Ser/Thr protein kinase, fibroblast growth factor receptor and regulation of endothelial cell migration and sprouting angiogenesis.

**Figure 6:**
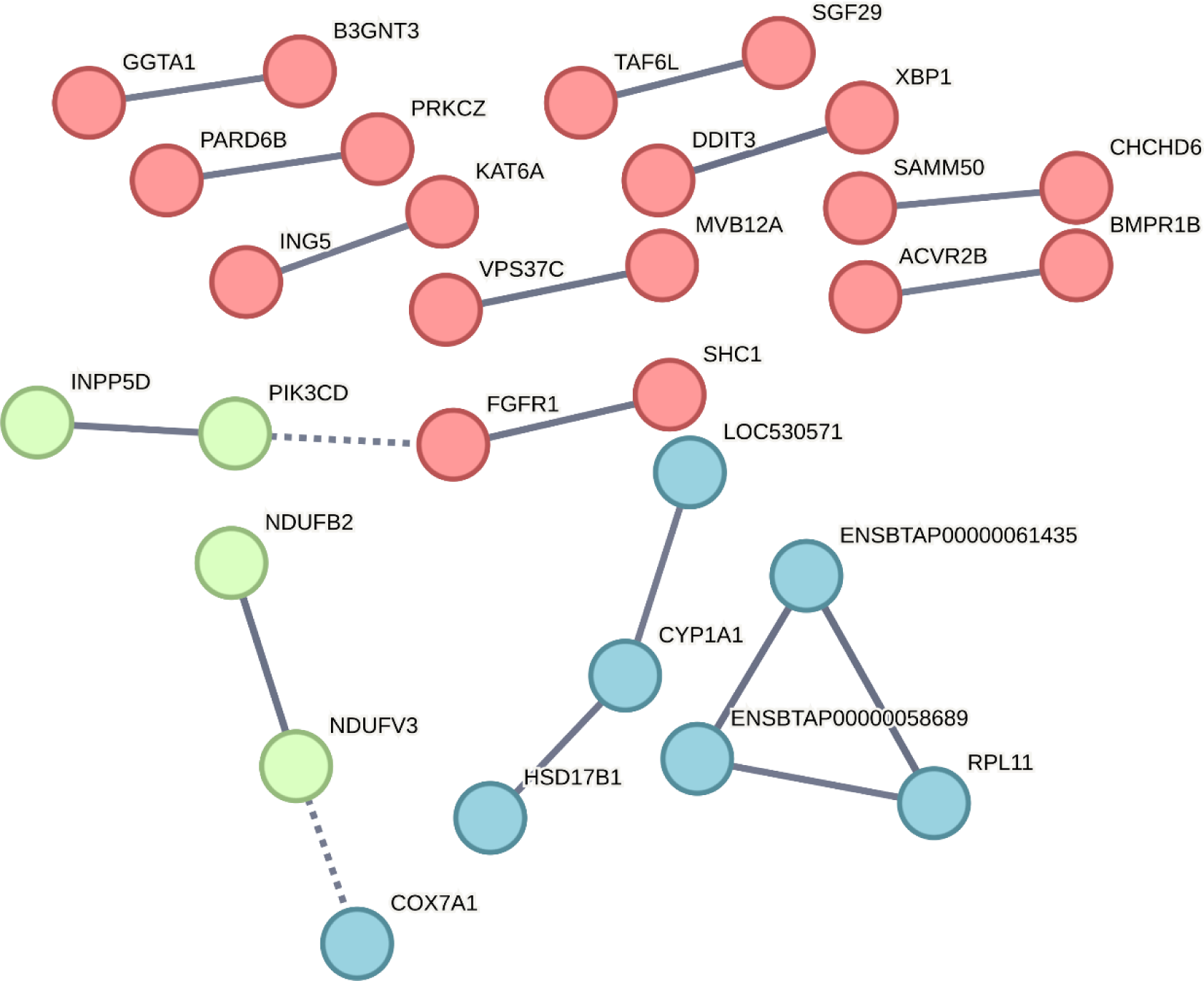
String protein interaction network of identified genes with differential methylation between healthy and diseased calves within 10kb of a TSS.

Extension of differential methylation analysis to base level identification of differential methylation in promotor and exon regions identified 4763 genes with differential methylation in their promotor regions and 65 in their exon regions. Panther overrepresentation test revealed 23 enriched GO biological process’, shown in Table 4, and three enriched Reactome pathways: protein-protein interactions at synapses (fold enrichment=2.86, p=4.77E-05, FDR=2.04E-02), neuronal system (fold enrichment =2.01, p=1.21E-08, FDR=6.87E-06) and signal transduction (fold enrichment =1.35, p=4.19E-09, FDR=3.58E-06). No functional enrichment was observed for genes with differential methylation in exon regions.

**Table 4:** The enriched biological process’ identified for the genes with differential methylation between healthy and BRD calves identified in their promotor regions as determined by PANTHER Overrepresentation Test Fisher Exact test.

| GO biological process | Fold Enrichment | P-value | FDR |
| --- | --- | --- | --- |
| biological regulation (GO:0065007) | 0.79 | 3.15E-06 | 2.27E-03 |
| regulation of biological process (GO:0050789) | 0.79 | 4.75E-06 | 3.02E-03 |
| response to stimulus (GO:0050896) | 0.76 | 1.81E-05 | 9.78E-03 |
| cellular process (GO:0009987) | 0.72 | 4.84E-15 | 1.75E-11 |
| biological_process (GO:0008150) | 0.7 | 2.25E-20 | 1.22E-16 |
| cellular metabolic process (GO:0044237) | 0.69 | 2.46E-08 | 2.96E-05 |
| organonitrogen compound metabolic process (GO:1901564) | 0.66 | 1.13E-07 | 1.11E-04 |
| protein metabolic process (GO:0019538) | 0.66 | 5.46E-06 | 3.28E-03 |
| primary metabolic process (GO:0044238) | 0.66 | 3.02E-11 | 5.44E-08 |
| metabolic process (GO:0008152) | 0.66 | 4.22E-13 | 1.14E-09 |
| organic substance metabolic process (GO:0071704) | 0.66 | 2.83E-12 | 6.12E-09 |
| nitrogen compound metabolic process (GO:0006807) | 0.65 | 7.15E-11 | 1.11E-07 |
| biosynthetic process (GO:0009058) | 0.65 | 2.25E-05 | 1.16E-02 |
| cellular biosynthetic process (GO:0044249) | 0.64 | 8.25E-05 | 3.72E-02 |
| organic substance biosynthetic process (GO:1901576) | 0.64 | 2.46E-05 | 1.21E-02 |
| organic cyclic compound metabolic process (GO:1901360) | 0.64 | 4.47E-06 | 3.02E-03 |
| macromolecule metabolic process (GO:0043170) | 0.64 | 4.78E-10 | 6.46E-07 |
| cellular aromatic compound metabolic process (GO:0006725) | 0.62 | 2.49E-06 | 1.92E-03 |
| cellular nitrogen compound metabolic process (GO:0034641) | 0.6 | 8.09E-08 | 8.76E-05 |
| heterocycle metabolic process (GO:0046483) | 0.59 | 5.38E-07 | 4.85E-04 |
| nucleobase-containing compound metabolic process (GO:0006139) | 0.59 | 1.13E-06 | 9.42E-04 |
| nucleic acid metabolic process (GO:0090304) | 0.59 | 1.29E-05 | 7.34E-03 |
| macromolecule biosynthetic process (GO:0009059) | 0.56 | 7.62E-05 | 3.58E-02 |

## Discussion

We set out this study to identify the differences in methylation profiles of healthy adults and calves and between healthy calves and those diagnosed with BRD using RRBS for the purpose of identification of markers of genetic age and of potential targets for genetic improvement of dairy cattle. Methylation differences between healthy calves and cows were primarily in regions associated with genes involved in growth and developmental pathways, as to be expected. Between healthy and diseased calves the identified differences in methylation were primarily in regions associated with genes involved in immune regulation and pulmonary pathways. These identified genes and regions potentially play a role insusceptibility to BRD and, therefore, may be able to be used to inform breeding programs to improve future herd health.

A higher percentage of hypomethylation observed in adult cows compared to calves is consistent with the established theory of genomic hypomethylation associated with ageing [58, 59], the small percentage however is likely associated with the small number of animals included in the study and the relatively young age of the adults (5-6 years old). Further the identification of differential methylation and enrichment of AKT Serine/Threonine Kinase 2 (*AKT2*) pathways between healthy cows and calves is an intriguing finding due to the implicated role of *AKT2* in aging. Murine experimental studies found *AKT2* ablation resulted in prolonged life-span with observed reduction in negative cardiac remodelling and a specific reduction in Ca2+ defects in contractile and intracellular cardiac tissue and mitochondrial injury, both typically associated with age [60]. Differential methylation affecting the regulation of this gene could therefore act as an important marker to establishing genetic ageing. It is, however, important to also note the role of *AKT2* in the immune system. *AKT2* is involved in macrophage polarization (ablation results in M2 macrophages involved in wound healing), regulation of the functions of dendritic cells and proliferation of T regulatory cells [61]. Due to the young age of calves included in the comparison, the enrichment of this pathway could also be indicative of age-associated immune system development.

Previous studies of differential methylation in aging have led to the production of epigenetic clocks, for establishing an organisms biological age [8, 62, 63], and identification of age related biological process with differential methylation [64]. In beef cattle, age related hypermethylation was observed in promoter and 5’ UTR regions, with hypomethylation in other regions. The observed hypermethylation was also found to be functionally linked to pathways including: DNA and transcription factor binding and regulation of genes associated with development and morphogenesis pathways and metabolism [63]. Similarly, previous functional enrichment analyses of hypermethylated genes in age comparisons of mice identified enrichment for functions such as system development, anatomical structure development, nervous system development, multicellular organismal development, cell fate commitment, cell differentiation, cell development, developmental process, cellular developmental process and multicellular organismal process [64]. In our study GO enrichment analysis of genes with methylation differences between healthy cows and calves were primarily involved in developmental pathways, for example tissue morphogenesis, neuron differentiation, regulation of developmental process, cell fate commitment and organ morphogenesis, consistent with previous findings [63, 64]. These enriched GO terms are expected as calves are still developing into adulthood, therefore, differential methylation of genes involved in these pathways could be useful candidates as markers of epigenetic ageing. Interestingly a reduction of GO enrichment of detection of chemical stimulus involved in sensory perception of smell was observed between healthy cows and calves and between healthy calves and those diagnosed with BRD. Many genes attributed in this GO annotation code for G protein coupled receptors, involved in cell communication and receptors to a range of stimuli, suggesting potentially enrichment of a more general signalling function outside of the olfactory system.

Resistance to BRD in dairy calves is a complex heritable trait [37, 38, 42], and therefore identification of genetic loci and candidate genes associated with susceptibility to BRD provide the opportunity for genomic selection aiming to reduce disease incidence. Nevertheless, epigenetic changes may also play a role since a significant number of genes have been identified in this study as differentially methylated between healthy calves and those diagnosed with BRD. Although we have not generated RNA-Sequencing data from these calves to compare their blood transcriptomic profiles, when we compared our results with the results of previous transcriptomics, eQTL and genome-wide association (GWAS) studies many of the differentially methylated genes have been also identified and prioritised as good candidate genes for BRD resistance in these studies [43, 65]. Specifically, eight genes (*KPNA6, NEIL3, ADCK1, ZNF507, REEP3, AHSA1, SEPTIN11* and IgLON5) identified as differentially methylated were also identified within significant GWAS windows in the study of BRD susceptibility in pre-weaned Holstein calves [65]. Further, some of the differentially methylated genes (*TGM3, NRG1, RETN, IL1R2, ADGRE1, ALPL, KREMEN1, TRPC5, MN1, ALOX5AP, DYSF, ITGA9, NUPR1, SLC28A3, IGLON5, PTPN5, GLT1D1, DPYS, WIPI1, CFB, LOC511106, IL3RA*) also overlapped with previously identified differentially expressed genes in studies of BRD susceptibility of pre-weaned calves and feedlot cattle [43, 65]. Of these, *NUPR1, CFB and ADGRE1* were also associated with identified trans-eQTLs [43].

Moreover, our study revealed that inflammatory response, pulmonary epithelial integrity, host innate and adaptive immune system regulation were significantly enriched pathways, similar as to what was reported in two previously conducted enrichment analysis of SNP data (GSEA-SNP) and multi-omic (transcriptomic, genomic and metabolomic) studies of BRD resistance in crossbred or multi-breed beef cattle and Holstein calves [43, 66]. Indeed, a number of innate immune related differentially methylated genes overlapped with existing identified important genes in BRD through genetic and transcriptomic studies; *PTPN5* (involved in interleukin-37 signalling), *ILR3A* (Interleukin 3 receptor subunit alpha), *IL1R2* (Interleukin 1 receptor type 2), *CFB* (involved in activation of C3 and C5 and alternative complement activation), *IGLON5* (immunoglobulin family protein), suggesting that epigenetic changes at the methylation level may underly these gene expression differences. The enrichment of the innate immune related genes in particular are consistently identified as important, consistent with findings in pre-existing studies [43], likely because at the age of calves at the time of sampling the adaptive immune system would not have been fully matured, unlike the innate immune system.

In particular, this comparison highlighted the enrichment of cardiac muscle contraction, regulation of the MAPK cascade, B cell differentiation and stress fibre assembly. Further protein network analysis revealed important differentially methylated genes and pathways related to pulmonary vasculature and respiratory cilia structure and function regulation providing further insights in the underlying mechanisms of BRD resistance in calves. A primary example of a good candidate gene is dynein axonemal intermediate chain 1 (*DNAI1*), part of the dynein complex found in respiratory cilia with a function in humans in regulating dynein activity, the means in which the motion and therefore function of the cilia is achieved [67, 68]. Moreover, genes with reported functions in inflammation pathways and in pulmonary vasculature are good candidates since in respiratory disease pulmonary vascular endothelial cells are among the primary cells to be damaged. Furthermore, platelet endothelial cell adhesion (*PECAM1*), is an adhesion molecule that facilitates leukocyte trans-endothelial migration during inflammation and is expressed on the surface of pulmonary vascular endothelial cells [69] and also interacts with the von Willebrand factor (*VWF*) which functions by promoting platelet adhesion to sites of vascular injury [70].

Identification of differential methylation at base level resolution applied to promotor and exon regions revealed a significantly high number of bases as differentially methylated in the promotor region of the gene that encodes CUB domain containing protein 2. CUB domain containing proteins are highly conserved with a diverse range of functions, for example: complement activation, inflammation, tissue repair and angiogenesis to name a few. In human studies, CUB domain containing protein (*CDCP1*) expression has been found to strongly correlate with poor prognosis and relapse of lung adenocarcinoma patients [71, 72]. The identification of the differentially methylated bases in this gene and its implication in human lung adenocarcinoma could be informative of a potential role in BRD. Significant hypomethylation in calves with BRD was also observed in the promotor region of *DNAI1* upon base level resolution analysis. A high number of bases were also identified as hypomethylated between healthy and BRD calves in exon regions of the endothelin converting enzyme 1 (*ECE1*) gene. This gene functions by converting big endothelin to biologically active peptides with roles in vasoconstriction [73, 74]. Increased expression of *ECE1* and its target endothelin-1 (*Et-1*) has previously been implicated in studies in pulmonary fibrosis in rat models [75–78] and in human idiopathic pulmonary fibrosis and correlated with disease activity [73, 78]. This, in combination, with the identified enrichment of circulatory system development and morphogenesis of epithelium is consistent with existing identified enrichment of pulmonary epithelial integrity [66] and is suggestive of an important mechanism in susceptibility to BRD.

In conclusion, functional enrichment of differentially methylated genes between healthy adult cows and healthy calves identified the expected developmental pathways with the exception of an enrichment of chemical stimulus involved in sensory perception of smell. We hypothesise however that this finding is due to the high number of G-protein coupled receptors assigned this GO annotation and that these receptors may have an alternative function in different tissues, currently as of yet unannotated. The identification of a high number of differentially methylated immune related genes between healthy and BRD calves in this study and the observed overlap of genes with those previously identified by GWAS, transcriptomic and eQTL studies reinforce the biological importance of these candidate genes and pathways. Based on the findings here, regulation of key genes such as *DNAI1, CDCP1* and *ECE1* may have roles in BRD resistance. In particular; due to the structural function in respiratory cilia, roles in vasoconstriction and existing implication in pulmonary fibrosis, *DNAI1* and *ECE1* respectively, are of great interest. Due to its role in complement activation and the existing GWAS and trans-eQTL evidence of the involvement of the complement cascade in BRD, we also consider *CDCP1* a key gene. It can, therefore, be concluded that resistance to BRD in calves is multifactorial with differential regulation in genes in immune, inflammatory and pulmonary vasculature significantly contributing to calf health with respect to BRD.

## Methods

### Ethical approval

This study was performed under the ethical approval of the University of Liverpool Research Ethics Committee (VREC927). Procedures regulated by the Animals Scientific Procedures Act were operated under Home Office License (P191F589B).

### Study Population

This study prospectively enrolled calves at one commercial dairy farm in North Wales. The farm was chosen due to its proximity to the University of Liverpool, and their willingness to participate. Calves were enrolled within one week of birth and were eligible for enrolment if Holstein, female, and were 1 to 7 days old when the weekly visit occurred. A full clinical examination was conducted at <1 week of age (“neo-natal”), 6 weeks of age (“pre-weaning”) and 15 weeks of age (“post-weaning”). Calves were housed individually in pens for the first two weeks of life in a farm building dedicated to housing neonatal calves. Thereafter two to three calves were grouped together in pens which had an outside feeding area and a covered deep straw bedded calf hutch. Calves were fed 4 L of colostrum in their first 4-6 h of life using a bucket and teat system, and then placed onto a commercially available milk replacer. Calf starter (concentrates) was available to all calves once moved into grouped housing.

Samples were also collected from adult healthy cows (Holstein, female, 5-6 years old) raised on the same farm. These animals were sampled as part of a routine herd health screening of freshly calved cows and were clinically healthy at the time of sampling.

### Data Collection

At enrolment (within one week of birth), a blood sample was collected. A full clinical exam was performed by a qualified veterinarian. Animals were also assessed using the Wisconsin scoring system [44] and a composite health scoring system. This scoring system devised by the researchers and is described in supplementary Table S2. Briefly, calves were given clinical examinations at weeks one, five and eight of the study and the measures listed in the table assessed along with record of any discharges associated with eyes and nose, ear position, coughing frequency and whether scour present. Once enrolled, calves were assessed weekly using the Wisconsin scoring system (scores taken for discharges associate with eyes and nose, ear position, coughing frequency and whether scour present) for the presence of bovine respiratory disease. At one, five, and eight weeks, calves underwent a full clinical examination and the Wisconsin scoring system and composite scoring system was used to assess health status. At eight weeks old, in addition to those measures already taken, thoracic ultrasound scanning [79] was also performed to assess respiratory health. Briefly, lungs were examined via ultrasonography and scored on a 0-5 system [79]. Score zero would be considered normal, score one indicates some damage but would not be considered significant. Score two and upwards would be considered significant lung damage. Score two is indicative of a patchy pneumonia/lung damage, whilst score three indicates significant lung damage and subsequent consolidation of a whole lung lobe. Scores four and five are as score three but indicate how many lung lobes are severely affected, such that score four affects two lung lobes and score five affects three or more lung lobes.

The three calves designated healthy control animals were calves 1-3 with no observed incidence of respiratory disease over the study period. The other three calves were diagnosed with BRD based on the above-described assessment.

### RRBS sequencing

DNA was extracted from whole blood samples from six calves (three healthy and three diseased) and four adult cows using the DNEasy Kit (Qiagen) according to the manufacturer instructions.

RRBS library preparation and Sequencing (paired-end 50 bp reads on a NovaSeq 6000 platform) was outsourced to Diagenode ((https://www.diagenode.com/en) using Illumina technology.

### RRBS sequencing analysis

Initially, FastQC (Andrews, 2010) version 0.11.8 was used for quality control of sequencing reads. Then, the software Trim Galore! Version 0.4.1 (Krueger, 2015) was used to remove sequencing adapters. Subsequently, Bismark version 0.20.0 (Krueger and Andrews, 2011), a specialised tool for mapping bisulfite-treated reads, was used to align reads to the reference genome (bostau8). Bismark requires that the reference genome, in this case bostau8 (source required), first undergoes *in silico* bisulfite conversion and transformation of the genome into forward (C - > T) and reverse strand (G -> A).

Reads producing a unique best hit to one of the bisulfite genomewere compared to the non-bisulfite converted genome to identify cytosine contexts (CpG, CHG or CHH – where H is A, C or T). The modules of Bismark version 0.20.0 (Krueger and Andrews, 2011): “cytosine2coverage” and “bismark_methylation_extractor”, were used to infer all cytosine methylation states, their context and to calculate percentage methylation. Only CpGs present in all samples were retained for further analysis.

DNA methylation analyses were conducted using methylKit version 1.20.0. [80]. The cytosine2coverage files output from Bismark were used as the methylation call files to create a methylRaw object. Descriptive statistics were calculated, and samples filtered based on read coverage (bases having lower coverage than 10 discarded and bases having higher coverage than 99.9% discarded). Samples were then merged by case/control (diseased/healthy and calf/cow) and correlation between them calculated before identification of differential methylation. Differential methylation in 1000bp regions using 500bp steps was identified using Chi-squared test with Sliding Linear Model (SLIM) correction for multiple testing. Differential methylation was determined based on a ≥25% difference in methylation and a statistically significant p value of ≤0.05 and q ≤0.01 between sample groups [80–82]. Further, differential methylation at base level resolution was identified, also based on a 25% difference in methylation and a statistically significant p value of ≤0.05 and q ≤0.01 between sample groups.

Differential methylation analyses, as described above, were conducted between healthy adult cows and healthy calves, healthy calves and those diagnosed with BRD (diseased) and finally between a set of twin calves, one healthy and one diseased.

### Functional enrichment and protein interaction network analyses

Genes with associated differentially methylated bases within 50kb of the transcription start site (TSS) were analysed for gene ontology (GO) term over and under representation to increase understanding of biological pathways and functions that are differentially methylated between healthy cows and calves as well as healthy and diseased calves. For the twin comparison due to the high number of identified genes only those within 1kb of the TSS were analysed for GO term enrichment. GO term over and under representation was assessed with DAVID functional enrichment analysis and PANTHER (v17.0) [83] overrepresentation test. Pathway enrichment was also assessed using Reactome v. 77 [84] against the Bos Taurus genome (Ensembl release 108) with calculation of the false discovery rate.

Protein interaction networks for differentially methylated regions within 1kb of a TSS in each comparison were analysed using STRING (v 11.5) [55] with parameters: minimum required interaction score - highest confidence (0.900), no second shell interactions, network edges - confidence, disconnected nodes hidden, kmeans clustering of interactions into 3 clusters.

## Acknowledgements

The authors acknowledge the financial support provided by the Biotechnology and Biological Sciences Research Council (BB/S002960/1, BB/S003614/1, BB/S002944/1)

The authors express gratitude to the farm staff and herd managers for their assistance and cooperation without which this study would not have been possible.

## Data availability

Raw data and methylation call files are available at ENA under the project accession PRJEB71365.

## Author contributions

AP, GO and GB: conceived and designed the study and secured funding.

EA performed the RRBS analysis and wrote the manuscript with input from all authors.

BG and GO collected the samples and performed the phenotyping.

AP extracted DNA and supervised the RRBS analysis with input from DX and DW.

DX provided expertise on enrichment analyses and DW expertise on BRD immunological responses.

All authors contributed to the interpretation of the results and assisted in revising the manuscript.

