## Supplementary data for "Identification of methylation markers for age and Bovine Respiratory Disease in dairy cattle"

*Supplementary Table S1: Dam and Sire information of the six calves included in the epigenetic analysis demonstrating their relatedness.*

| Calf | Dam | Sire |
|---|---|---|
| Calf 1 | 3608 | INVICTUS |
| Calf 2 | 2861 | HO5593 CROSBY |
| Calf 3 | 2931 | HO5593 CROSBY |
| Calf 10 | 3581 | INVICTUS |
| Calf 11 | 2829 | HO5593 CROSBY |
| Calf 12 | 2931 | HO5593 CROSBY |

*Supplementary Table S2: Composite health scoring system used to diagnose Bovine Respiratory Disease in calves.*

| Measure | Score 0 | Score 1 | Score 2 |
|---|---|---|---|
| Demeanor | Normal-BAR | Dull/Listless (with slowed or staggered response to stimulus) | Moribund |
| Appetite | Good | Ok | Poor |
| Exercise Intolerance | None | Present | Marked |
| Teeth Grinding | None | Present Occasionally | Present Often |
| Clinical Exam (presence of injuries/body lesions) | Normal | Abnormal | Grossly Abnormal |
| Rumen Appearance | Hollow | Normal | Bloated |
| Signs of Pain (Bruxism, Abnormal Vocalisation, Abnormal Posture) | None | Occasional | Frequent |
| Hair Coat quality (Dull, Hairloss, Thicker coat) | Excellent | Ok | Poor |
| Cleanliness of Calf (Dirty flanks, Perineum, Presence of Diarrhoea) | None | Small amounts (less than a hand area) | Larger amounts (over a hands area) |
